# Chemokine-induced cell swelling fluidises the plasma membrane to regulate CD8^+^ T cell migration

**DOI:** 10.64898/2026.10.07.757276

**Authors:** Leonard L. de Boer, Cenk Onur Gurdap, Erdinc Sezgin

## Abstract

T cell migration is primarily viewed as a cytoskeletal process. Here, we show that chemokine-stimulated CD8^+^ T cell migration is modulated by a volume-driven biophysical switch at the plasma membrane (PM). We found that chemokine-induced cell swelling creates an adhesion-dependent gradient of high PM fluidity at the leading edge. External chemical fluidisation of the PM using membrane modulators enhances migration similar to chemokine-induced fluidisation without the need of cell-swelling. Together, our findings identify PM fluidity as a tuneable biophysical gatekeeper that controls T cell migration, potentially providing a novel mechanical tool for optimising adoptive cell therapies in restrictive tissue microenvironments.

**One sentence summary:** Chemokine-induced cell swelling increases plasma membrane fluidity at the leading edge to support actin retrograde flow dynamics.

**Significance Statement:** Immune cells must move rapidly through the body to find and eliminate threats, but how they physically control movement has remained unclear. We show that chemokine signals trigger T cells to take up water and swell, causing their plasma membrane to become more fluid. This local membrane fluidization reduces mechanical resistance at the cell front, enabling the actin cytoskeleton to push the membrane forward and drive migration. Importantly, cell adhesion concentrates this fluid state at the leading edge, creating a spatial cue that directs movement. Our findings identify the plasma membrane as an active mechanical regulator of cell migration and provide a new framework for understanding immune-cell behavior, cancer-cell invasion, and the design of more effective cell-based immunotherapies.

## Introduction

Migration is a crucial part of T cell function. Naïve CD8^+^ T cells constantly recirculate and migrate within the bloodstream and dense lymphatic tissues in search of cognate antigen presented on antigen-presenting cells, an essential task for initiating adaptive immunity (*1*). T cells are guided to antigen-presenting cells by the chemokine CCL19, secreted by stromal and dendritic cells. CCL19 stimulation triggers large-scale changes in T cell shape and function, polarising the cell and initiating migration (*2*). At the front, or leading edge, of the migrating T cell, large-scale reorganisation of the cytoskeleton drives the assembly of a dense, branched F-actin network that physically pushes the plasma membrane (PM) forward (*3*). At the nanometre scale, this protrusive force can be described by the Brownian ratchet model, where stochastic thermal fluctuations of the lipid bilayer and the actin filament create transient gaps into which G-actin monomers insert and attach to the filament, thereby protruding the membrane forward (*4, 5*).

While migration has classically been viewed as a purely cytoskeleton-driven process, new work has shown that T cell migration and activation require a coordinated and localised ion and water influx (*6*–*8*). Chemokine signalling activates inward Na^+^ and K^+^ pumps, driving an osmotic water flux that increases T cell volume by 10-20% (*7*). Genetic or pharmacological disruption of swelling strongly impacts migration speed and limits actin polymerisation rates across multiple immune cell types (*7, 9*–*11*). These observations link intracellular water flux and the actin cytoskeleton. Crucially, however, how these volume changes alter the highly ordered physical state of the T cell PM (*12*) remains poorly understood. To accommodate volume expansion and thereby prevent cell rupture (*13*), T cells possess a large surplus of membrane stored in highly folded reservoirs (*14*). While unfolding these reservoirs has been shown to expand cell surface area (*14*), the biophysical impact of this has not been explored. Thermodynamically, membrane curvature and lipid packing are linked: tightly curved membrane folds restrict the area per lipid headgroup, thereby compressing the outer leaflet and thus reducing lateral lipid motility (*15, 16*). Conversely, flattening or smoothing out these folds by expanding cell volume should reduce the mechanical compression and thus result in an increase in membrane fluidity. A local increase in hydraulic pressure, aside from smoothing membranes, can also alter membrane-cytoskeleton tethering. Breaking physical attachments between the PM and the actin cytoskeleton has been shown to be required for protrusion initiation (*17*), and, somewhat counterintuitively, F actin is depleted proximal to the PM (*7, 18*). Because the cytoplasm behaves like a poroelastic sponge, preventing instantaneous equilibration of water in the cell (*19*), a directional water influx can focus hydrodynamic pressure to help facilitate actin-PM uncoupling (*7, 20*). Removing the actin scaffold anchoring to the PM removes a significant barrier to lipid mobility(*21*–*23*).

Recent work has established that membranes can sustain local and stable tension gradients across distinct zones rather than equilibrating them instantaneously across the entire cell surface (*24*). This mechanical compartmentalisation suggests that local changes in membrane fluidity can also be compartmentalised in a similar way. Local changes in fluidity could increase the gap-forming rate required for the Brownian ratchet driven actin filament elongation. In this study, we use the environment-sensitive probe NR12A to determine the spatiotemporal dynamics of PM fluidity in migrating human naïve CD8^+^ T cells. We show that chemokine stimulation triggers a rapid increase in PM fluidity that is coupled to an increase in cell size. We show that this PM fluidity shift is spatially organised, forming a non-linear fluidity gradient from the more fluid leading edge to the less fluid trailing edge in adherent cells. Finally, by decoupling cell volume changes from PM lipid packing, we demonstrate that external chemical PM fluidisation rescues actin retrograde flow velocities and migration speeds in non-swelling cells. Taken together, our study demonstrates that PM fluidity is a tuneable biophysical parameter that modulates T cell migration and coordinates actin cytoskeletal dynamics.

## Results

### CCL19 stimulation induces T cell PM fluidisation

To measure variations in membrane organisation, we used the solvatochromic probe NR12A (*25*), which monitors the extent of water penetration into the lipid bilayer through a solvent relaxation mechanism. In fluid or loosely packed environments, increased hydration around the fluorophore leads to a red shift in emission, providing a ratiometric readout of local lipid packing (Fig 1A). We imaged blood-derived primary human naïve CD8^+^ T labelled with NR12A migrating under agarose on dishes coated with ICAM-1 to mimic the mechanical confinement of the tissue microenvironment (*26*). Upon CCL19 stimulation, T cells rapidly initiated migration under agarose confinement (Fig 1B, supplementary movie 1). Ratiometric analysis of the NR12A signal revealed a global decrease in PM generalised polarisation (GP) from approximately 0.22 down to 0.12 (Fig 1C-D), indicating a robust reduction in lipid packing, corresponding to an increase in PM fluidity. We verified that this fluidisation was accompanied by chemokine-induced osmotic swelling as previously described (*7*–*9*), with modal cell volume increasing from a baseline of 170 fl to ∼200 fl following chemokine stimulation (Fig 1E).

**Figure 1:**
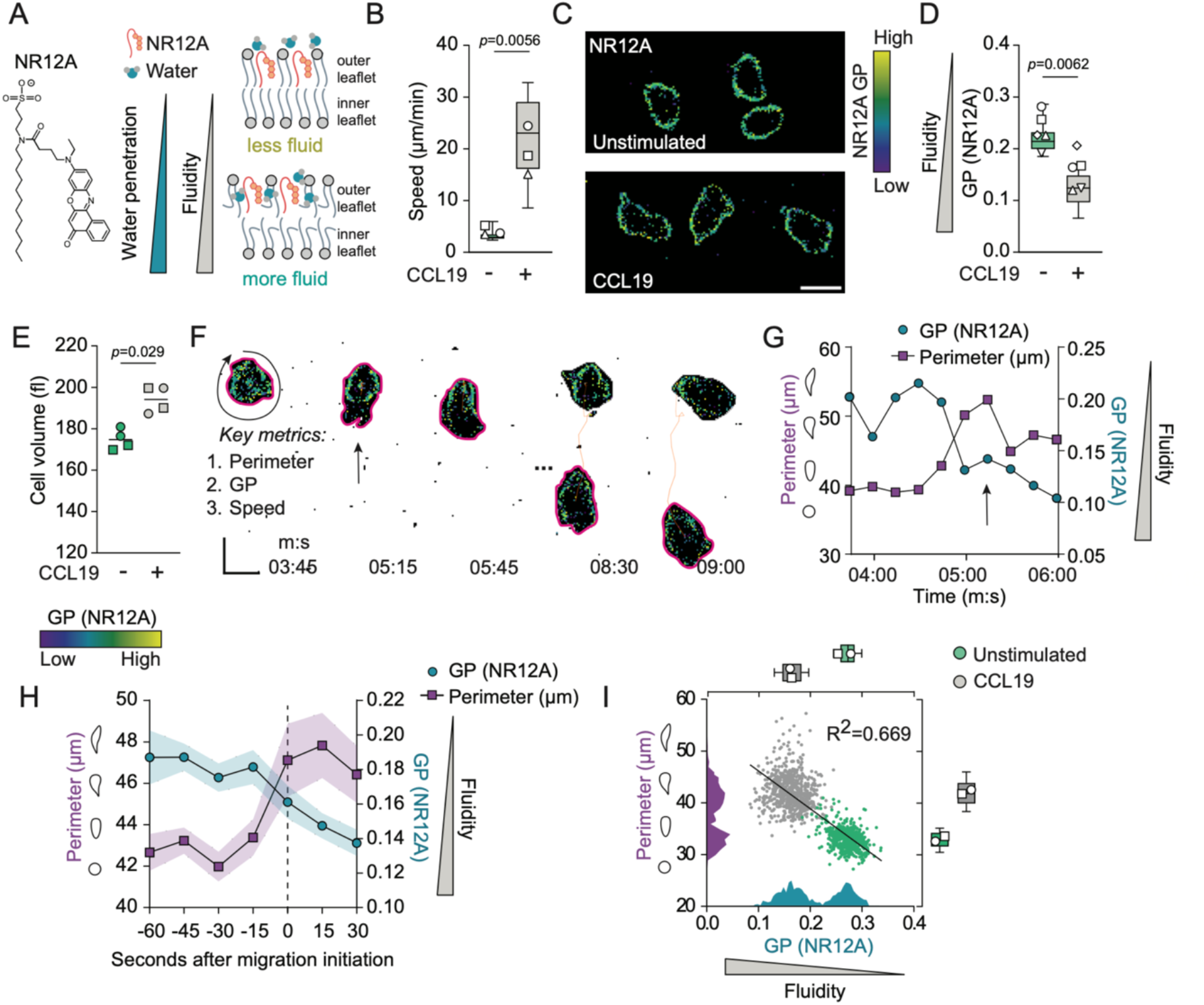
Rapid membrane fluidisation follows CCL19 stimulation of CD8^+^ T cells. **(A)** Schematic representation of the ratiometric solvatochromic probe NR12A. In fluid or loosely packed lipid environments, increased water penetration into the outer leaflet of the PM causes a red shift in fluorescence emission, yielding lower generalised polarisation (GP) values. In tightly packed environments, restricted water penetration in the outer leaflet of the PM causes a green-shifted emission, resulting in higher GP values. **(B)** Speed of naïve human CD8^+^ T cells migrating under agarose in response to CCL19. **(C)** Example images of the general polarisation (GP) signal of human naïve CD8+ T cells under agarose stained with NR12A. Cells were either unstimulated or stimulated with CCL19. **(D)** GP of individual cell shapes in naïve CD8+ T cells migrating response to CCL19. **(E)** Cell volumes of naïve CD8+ T cells stimulated with CCL19 for 20 min. Each dot represents the modal volume of ∼2000 cells determined using a Casyton. Lines represent means. **(F)** Example image of a naïve CD8+ T cell polarising and starting to migrate in response to CCL19 stimulation. Black arrow indicates migration initiation. Pink shapes indicate the TrackMate shape detection from which cell perimeter and speed are derived, and the GP measure is the average of the signal from within the cell shape. Orange lines represent the cell track over time. **(G)** Quantification of perimeter (purple) and GP (cyan) of the cell from F. Black arrow indicates migration initiation. **(H)** Mean (±SEM, shaded area) of perimeter and GP of cells before and after migration initiation (dashed line). **(I)** Perimeter of individual naïve CD8+ cell shapes relative to GP (data from 2 experiments with corresponding shape in D). Histograms represent frequency distribution (perimeter, purple; GP, cyan). Black line represents linear regression. R is correlation coefficient. GP, generalised polarisation. Sample numbers: Unstimulated *n =*218 tracks, CCL19 *n =* 296 tracks (B); unstimulated, *n=*3567 cell shapes; CCL19, *n=*5510 cell shapes (D); *n* = 4 biological replicates where each shape represents 1 experiment (E), *n* = 1 cell (G), *n =* 19 cells (H), unstimulated, *n=*586 cell shapes, CCL19, *n*=737 cell shapes (I). Scale bar, 5 µm (C,F). Data are pooled from 2 (E, I), 3 (B, H) or 5 (D) experiments. Each overlaid symbol represents the mean of one experiment. Individual shapes represent experiments: in E each corresponding shape shows the biological replicates in each experiment. Box-plot centre line, median; box limits, 75th and lower 25th percentiles; whiskers, 10th and 90th percentiles. Statistical analysis carried out using a nested T test (B, D), Mann-Whitney test (E) or Spearman’s correlation coefficient (I). GP, generalised polarisation.

We next evaluated the temporal relationship between cell morphology and PM fluidity to determine how cell spreading relates to local membrane packing. Tracking individual cells over time allowed us to correlate changes in 2D perimeter with PM GP (Fig 1F, supplementary movie 2). At the point of migration initiation, both cell perimeter and membrane fluidity exhibited a concurrent increase (Black arrow, Fig 1F, G). When tracking multiple cells, we observed that cell perimeter expanded prior to the start of migration, reflecting initial cell polarisation and elongation (Fig 1H). Following migration initiation, PM fluidity increased further.

To define the relationship between unstimulated and stimulated cell states, we correlated cell perimeter against GP for both conditions (Fig 1I). We observed a biphasic distribution, where unstimulated cells clustered as small, low-perimeter cells with a less fluid PM, whereas CCL19-stimulated cells shifted to a larger, more fluid state. Together, these results demonstrate that upon chemokine stimulation, T cells undergo polarisation, swelling and elongation, which is followed by a progressive fluidisation of the PM.

### Localised PM fluidisation requires substrate adhesion

Following the temporal characterisation of PM changes, we investigated its contextual and spatial dynamics across different microenvironments, namely in suspension and in non-attached agarose confinement. Stimulating CD8^+^ T cells in suspension revealed that average PM GP did not shift in a consistent direction upon chemokine exposure (Fig 2A). We next evaluated whether physical confinement under agarose induces PM fluidisation in the absence of substrate adhesion. When chemokine-stimulated T cells were placed on a non-adherent, poly-ethylene glycol (PEG)-coated surface, the global drop in PM GP was notably less compared to ICAM-1 attachment (Fig 2B), indicating that global lipid packing shifts under confinement are modest in the absence of substrate attachment. We therefore next addressed the role of active cellular attachment on chemokine-induced biophysical remodelling.

**Figure 2:**
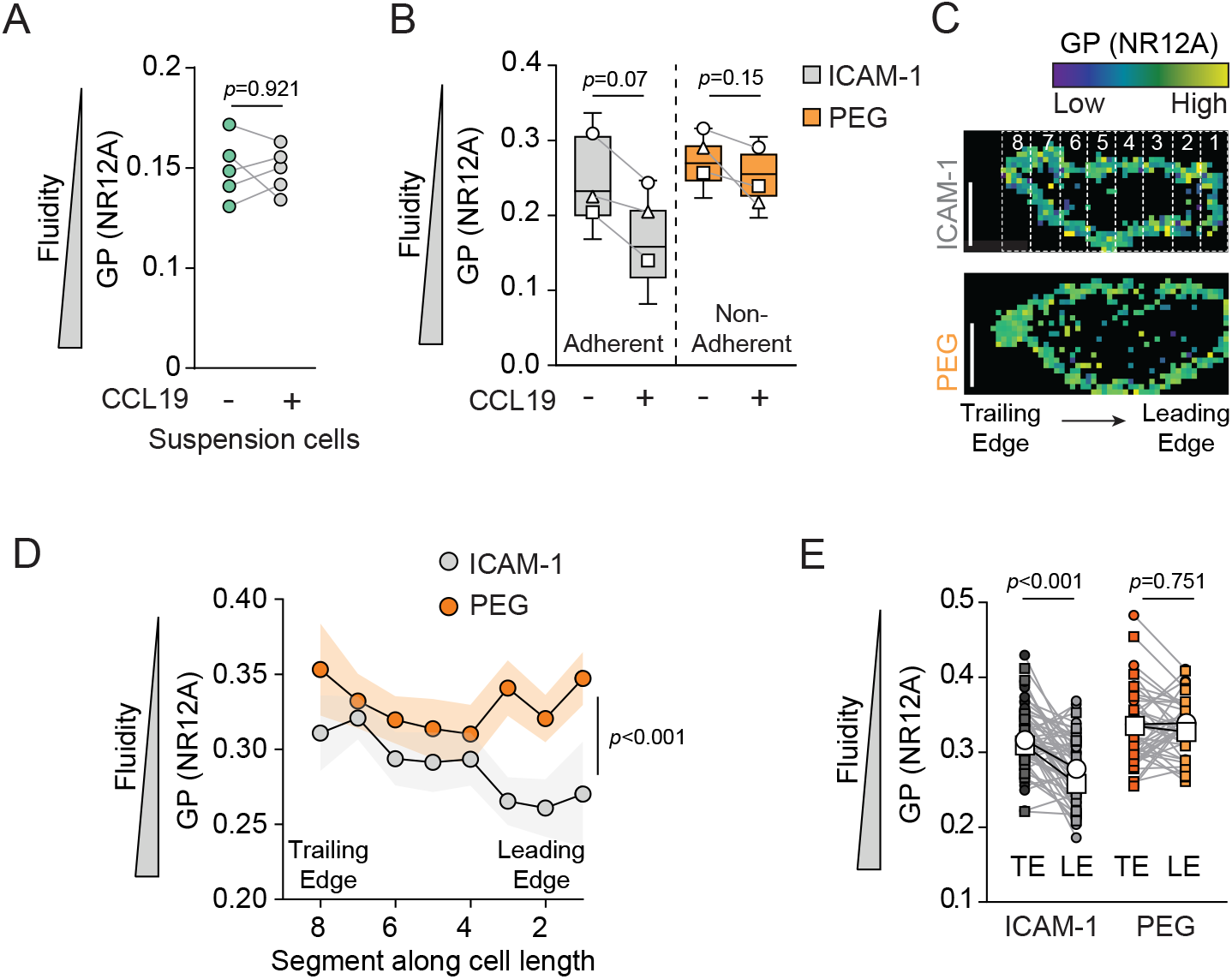
Membrane fluidisation is adhesion dependent and localised to the leading edge. **(A)** Percent change in GP of naïve CD8^+^ T cells in suspension following CCL19 stimulation for 10 min, measured by spectral flow cytometry. Dots represent T cells from individual donors. **(B)** GP of naïve CD8^+^ T cells under agarose, either unstimulated or stimulated with CCL19. Cells are either on a surface coated in ICAM-1 (adherent) or poly-ethylene glycol (PEG, non-adherent). **(C)** Example CCL19-stimulated naïve CD8^+^ T cells on either ICAM-1 or PEG under agarose, labelled with NR12A. Numbered white boxes represent segments used to generate data in D. Arrow indicates direction of migration or polarisation. **(D)** GP per cell segment along the longitudinal cell axis from the trailing edge (segment 8) to the leading edge (segment 1) of the cell. Dots represent mean GP per cell segment, shaded area represents SEM. **(E)** Paired comparison of the mean GP from the first two (1-2) (LE, leading edge) and the last (TE, trailing edge) two (7-8) segments per cell (same data as D). Coloured dots represent individual cells, lines connect segments from each cell. Sample numbers: *n*=5 biological replicates for each (A); unstimulated ICAM-1, *n* =385 cell shapes; CCL19 ICAM-1, *n*=1560 cell shapes, unstimulated PEG, *n*=2803 cell shapes, CCL19 PEG, *n*=2873 cell shapes (B); ICAM-1, *n* =55 cells, PEG, *n*=43 cells (D); same data as in D (E). Scale bar, 5 µm (C). Data are from 1 (A) or pooled from 2 representative of 3 (D,E) or pooled from 3 (B) experiments. Each overlaid shape represents the mean of one experiment, lines represent mean. Box-plot centre line, median; box limits, 75th and lower 25th percentiles; whiskers, 10th and 90th percentiles. Statistical analysis carried out using Mann-Whitney U-test (A), 1-way ANOVA (B,E), 2-way ANOVA (D). GP, generalised polarisation.

Active T cell protrusion depends on LFA-1 coupling ICAM-1 to the underlying actin cytoskeleton (*26*). Analysis of migrating cells revealed that on ICAM-1, PM fluidity is highly compartmentalised, demonstrating higher fluidity at the leading edge compared to the trailing edge (Fig 2C, D). The transition from the trailing edge to the fluidised leading edge was not linear, characterised by a step-drop in GP near the front third of migrating cells (Fig 2D). Conversely, on non-adherent PEG surfaces, this spatial gradient was completely absent, with the leading and trailing edges displaying similar fluidities (Fig 2E). Together, these data demonstrate that chemokine signalling alone is insufficient to drive localised PM biophysical changes, and that substrate adhesion is required to control strong spatial compartmentalisation of PM fluidity during migration.

### PM fluidisation state modulates T cell migration speed

Having observed the compartmentalized fluidity in the plasma membrane of stimulated cells, we next investigated the underlying mechanism driving these localised changes in PM fluidity. Because previous work has shown that the WNK1 pathway drives water influx at the leading edge of migrating cells (*7*), inhibiting WNK1 should prevent chemokine-induced T cell swelling.

We verified this by measuring T cell volume changes in response to CCL19 in the presence or absence of WNK463, a selective WNK inhibitor (*27*). Because a full inhibitory dose of WNK463 entirely prevents T cells from polarising and responding to stimuli (*7, 8*), we applied a low, partially inhibitory dose (0.4 µM) to restrict water entry while maintaining cellular responsiveness. We found that treating T cells with WNK463 reduced baseline cell volume and successfully blocked chemokine-induced swelling (Fig 3A). We next asked whether this reduction in cell volume altered membrane biophysical properties. Indeed, WNK463-treated cells showed a significant decrease in PM fluidity compared to vehicle-treated controls (Fig 3B), which was accompanied by a severe reduction in migration speed (Fig 3C).

**Figure 3:**
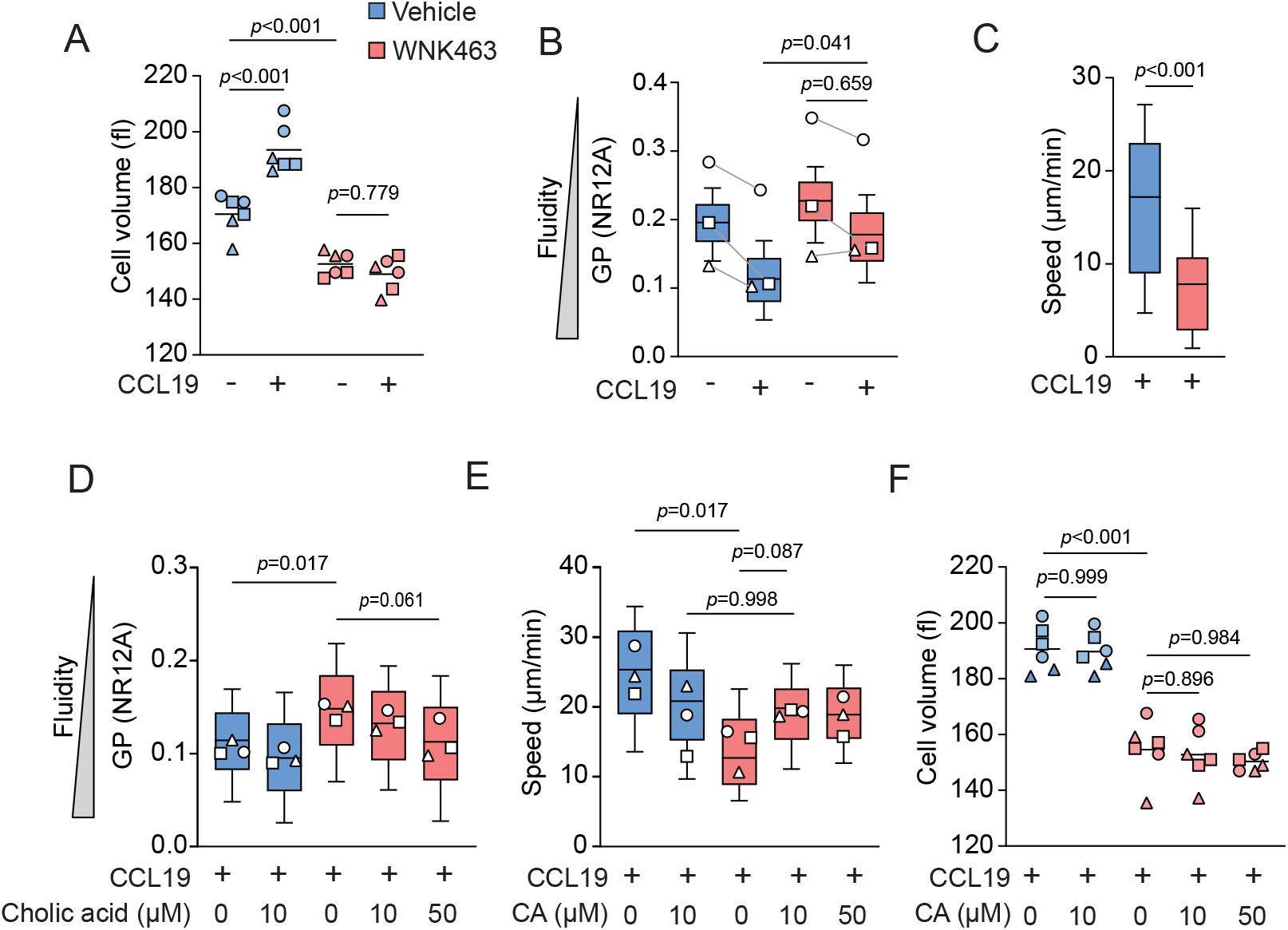
Membrane fluidisation promotes CD8 T cell migration. **A)** Cell volumes of naïve CD8^+^ T cells pretreated with or without WNK463 (0.4 µM) for 20 min, stimulated with CCL19 for 20 min. Each dot represents the modal volume of ∼2000 cells determined using a Casyton. Lines represent means. **(B)** GP of naïve CD8^+^ T cells treated with or without WNK463 on ICAM-1 under agarose, either unstimulated or stimulated with CCL19. **(C)** Speed of naïve CD8^+^ T cells treated with or without WNK463, stimulated with CCL19 migrating under agarose on ICAM-1 **(D)** PM GP of CCL19-stimulated naïve CD8^+^ T cells on ICAM-1 under agarose, treated with vehicle or WNK463 and supplemented with cholic acid (CA) at the indicated concentrations (10 µM or 50 µM) **(E)** Speeds of the cells in D **(F)** as in A, with the addition of the indicated concentration of CA during 20 min of WNK463 pretreatment. Sample numbers: *n* = 6 for all conditions (A, F), Unstimulated ICAM1, *n* = 2183 cell shapes, CCL19 ICAM1, *n* = 2699 cell shapes, unstimulated WNK463 ICAM1, *n* = 1837 cell shapes, CCL19 WNK463 ICAM1, *n* = 1784 cell shapes (B), vehicle, *n* = 46 tracks, WNK463, *n* = 38 tracks (C),CCL19, *n* = 3630 cell shapes, CCL19 CA10, *n* = 3010 cell shapes, CCL19 WNK463, *n* = 1514 cell shapes, CCL19 WNK463 CA10, *n* = 2882 cell shapes, CCL19 WNK463 CA50 *n* = 2978 cell shapes (D), CCL19, *n* = 349 tracks, CCL19 CA10, *n* = 189 tracks, CCL19 WNK463, *n* = 168 tracks, CCL19 WNK463 CA10, *n* = 258 tracks, CCL19 WNK463 CA50 *n* = 217 tracks (E). Data are from 1 experiment representative of 2 (C) or pooled from 3 (A, B, D-F) experiments. Each overlaid shape represents the mean of one experiment. Box-plot centre line, median; box limits, 75th and lower 25th percentiles; whiskers, 10th and 90th percentiles. Statistical analysis carried out using 1-way ANOVA (A-B, D-F), or a Mann-Whitney U-test (C). *p* > 0.05.

We subsequently wanted to determine whether membrane fluidisation is a consequence of cell shape changes, or a strict physical requirement for forward migration. To directly test whether restoring PM fluidity could rescue the migration defect independently of cell volume, we externally fluidised the PM using cholic acid (CA). CA incorporates into the outer leaflet of the PM, physically spacing out phospholipid headgroups to increase lateral mobility (*28*). Addition of 10 µM CA slightly increased PM fluidity in vehicle-treated T cells (Fig 3D). Crucially, in volume-restricted WNK463-treated cells, CA treatment fluidised the more packed PM, with 50 µM CA driving the greatest increase in fluidity (Fig 3D).

We next asked how this change in external fluidity impacts cell speed. While 10 µM CA slightly decreased migration speed in vehicle-treated control conditions, it introduced a trend towards partially rescuing the speed defect seen in WNK463-treated T cells (Fig 3E). Increasing the CA concentration to 50 µM provided no additive increase in speed (Fig 3E). Importantly, this partial rescue in migration speed occurred without altering cell volume, which remained completely suppressed by the WNK463 treatment (Fig 3F). These observations indicate that exogenous fluidisation of the PM can partially decouple migration velocity from active volume-expansion pathways, highlighting PM fluidity as an important biophysical parameter during T cell migration.

### Osmotic volume changes and cortical actin reorganisation control PM lipid packing

While external chemical fluidisation with CA allowed us to bypass swelling, we next wanted to understand the physiological limits of this pathway. Specifically, we asked how direct osmotic volume changes and the physical constraints imposed by the underlying actin cortex cooperatively set baseline PM fluidity. To change cell volume, we exposed naïve CD8^+^ T cells to varying agarose tonicities (hypertonic: 375 mOsm/L, isotonic: 300 mOsm/L, and hypotonic: 150 mOsm/L) (Fig 4A–C). As predicted by our model, a hypertonic environment caused a decrease in cell perimeter accompanied by a decrease in PM fluidity (Fig 4B, C). Conversely, swelling cells in a hypotonic environment slightly increased the cell perimeter and resulted in an increase in PM fluidity (Fig 4B, C).

**Figure 4:**
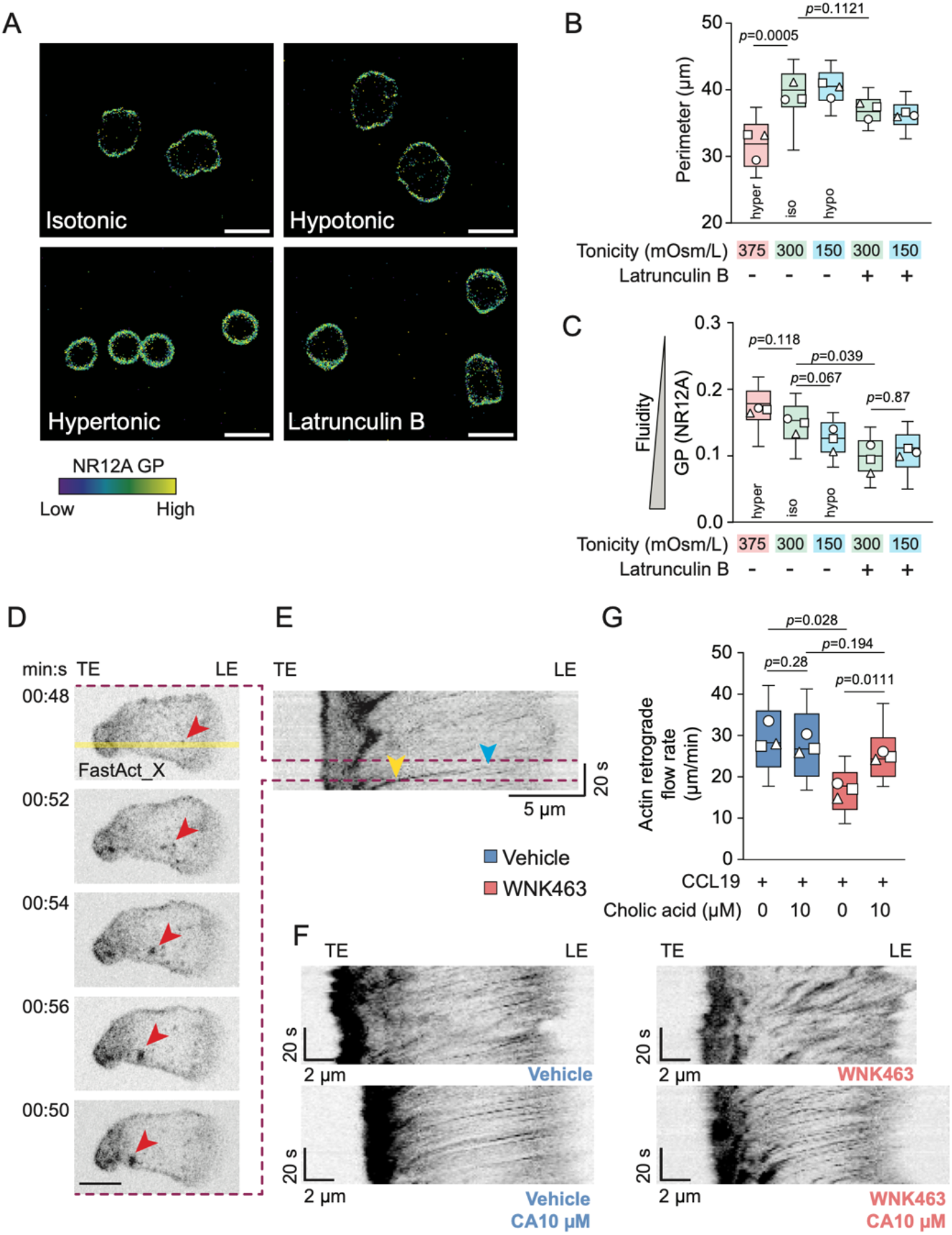
PM fluidity is dependent on cell volume and cytoskeleton release, and regulates actin polymerisation rates. **(A-C)** Naïve CD8^+^ T cells under either isotonic (iso), hypotonic (hypo) or hypertonic (hyper) agarose, with or without Latrunculin B. **(A)** Representative images of the membrane GP signal using NR12A under the indicated osmotic and inhibitor conditions. **(B)** Quantification of single cell perimeters across the indicated treatments to determine size changes. **(C)** Quantification of single cell PM GP across the indicated treatments to determine shifts in PM packing. **(D-G)** Naïve CD8 T cells stained with FastAct_X to visualise F-actin stimulated with CCL19 on PEG-coated dishes under agarose, imaged for 60s every sec. **(D)** Example of a cell showing retrograde flow of actin from the leading edge to the trailing edge. Red arrow indicates F-actin speck flowing rearward. Yellow line indicates the location of the kymograph in E. **(E)** Kymograph of the cell in D. Blue arrow indicates the start of the actin speck flow, yellow arrow indicates the end of the flow. Retrograde flow rates are calculated using the slope of line between the indicated start and end points (µm/min). Horizontal axis, distance, vertical axis, time. **(F)** Representative kymographs of F-actin flows in cells with the indicated inhibitors and cholic acid treatments. **(G)** Quantification of F-actin retrograde flow rates. Sample numbers: vehicle hypertonic *n* = 390 cells, vehicle isotonic *n* = 448 cells, vehicle hypotonic *n* = 392 cells, latrunculin B isotonic *n* = 524 cells, latrunculin B hypotonic *n* = 455 cells (B); vehicle hypertonic *n* = 364 cells, vehicle isotonic *n* = 430 cells, vehicle hypotonic *n* = 387 cells, latrunculin B isotonic *n* = 523 cells, latrunculin B hypotonic *n* = 439 cells (C); vehicle control *n* = 67 cells, vehicle + 10 µM CA *n* = 63 cells, WNK463 *n*= 59 cells, WNK463 + 10 µM CA *n*= 60 cells (G). Scale bar, 5 µm (A, D). Data are pooled from 3 experiments (B, C, G). Each overlaid shape represents the mean of one experiment. Box-plot centre line, median; box limits, 75th and lower 25th percentiles; whiskers, 10th and 90th percentiles. Statistical analysis carried out using a nested 1-way ANOVA (B,C,G).

Actin cytoskeleton is a key player for migration and PM is structurally supported by the actin cytoskeleton. Therefore, we next investigated the relationship between PM fluidity and the underlying actin network. We treated cells with latrunculin B, a small molecule inhibitor that binds G-actin monomers to prevent actin filament assembly. We observed that disassembling the cortical actin network under isotonic (300 mOsm/L) conditions caused an increase in PM fluidity (Fig 4C), matching the fluidisation response seen in hypotonic cell swelling. We next asked whether there would be additive PM fluidisation if we pushed these actin-disrupted cells further using a hypotonic environment. Surprisingly, exposing latrunculin B-treated cells to a hypotonic environment (150 mOsm/L) yielded no further increase in PM fluidity, with GP values remaining identical to the actin-disrupted isotonic condition (Fig 4C). Notably, we observed that these actin-inhibited cells failed to increase their perimeter when exposed to the hypotonic challenge (Fig 4B). These results imply that hypertonic cell shrinkage increases membrane lipid packing, whereas disassembling the cortical actin network reduces lipid packing to a minimum baseline threshold that cannot be lowered further by osmotic swelling. This indicates that the physical constraints imposed by the cortical cytoskeleton play a central role in modulating plasma membrane lipid organisation. Importantly, these findings raise a reciprocal question: if the actin cytoskeleton constrains PM fluidity, does the fluidisation state of the PM conversely act as a mechanical feedback barrier that tunes the rate of actin assembly?

### Membrane fluidity correlates with actin retrograde flow rate

To directly test whether PM fluidity correlates with cytoskeletal assembly rates, we evaluated actin polymerisation dynamics. T cell migration requires continuous actin polymerisation in the direction of migration (*3*). Membrane protrusion is driven by the addition of G-actin monomers to the barbed ends of F-actin filaments directly proximal to the PM (*29*). The Brownian ratchet model describes that thermal fluctuations of both the F-actin filament and the bilayer create transient stochastic gaps that allow G-actin monomers to insert and bind the F-actin filament (*5*). The filament consequently elongates, incrementally pushing the PM forward (*5*). We hypothesised that a more fluid lipid membrane facilitates filament growth. Because an increase in membrane fluidity reduces its local bending rigidity, it lowers the thermodynamic energy barrier required to deform the PM (28), thereby increasing the frequency and/or size of filament-membrane gaps where monomer insertion can occur. Conversely, when water influx is blocked in WNK463-treated cells, the PM remains locked in a highly ordered state (Fig 3B). This biophysical state could introduce a mechanical constraint on the underlying actin cytoskeleton, potentially restricting the dynamics of G-actin monomer addition and subsequent filament assembly at the membrane interface.

If membrane fluidisation regulates G-actin monomer addition, increasing fluidity in a packed membrane should thus accelerate retrograde flow rates. The retrograde flow of F-actin from the leading edge toward the cell body in the absence of substrate attachment serves as a readout of the rate of actin polymerisation (*26*). To evaluate this, we measured retrograde actin flow rates in CCL19-stimulated CD8^+^ T cells using the live-cell F-actin reporter SPY-FastAct. Cells were imaged under agarose on non-adhesive PEG-coated dishes. PEG prevents integrin-mediated gripping to the substrate, thus causing the polymerising actin network to slip backward rather than drive forward cell movement (supplementary movie 3). Kymograph analysis of actin flows revealed rapid retrograde flow in vehicle-treated control T cells (Fig 4D–F). Consistent with our hypothesis, flow rates were strongly reduced in volume-restricted, WNK463-treated cells (Fig 4F, G). Remarkably, supplementing WNK463-inhibited cells with 10 µM CA rescued actin retrograde flow rates back to control levels (Fig 4G). Notably, this rescue of the actin retrograde flow rate occurred despite cell volume remaining fully suppressed as measured previously (Fig 3F).

Taken together, these results show that PM lipid packing states are coupled to cortical actin dynamics, demonstrating that chemical fluidisation of the PM can bypass volume-restriction blocks to restore actin retrograde flow rates.

## Discussion

In this study, we show that chemokine signalling in CD8^+^ T cells drives cell polarisation and swelling, which directly increases PM fluidity. Notably, we demonstrated that PM fluidisation is an important requirement for both cell migration and, critically, for regulating actin dynamics.

To explain this, we propose a unified biophysical model where chemokine stimulation initiates a mechanical chain reaction: WNK1-dependent cell swelling increases the average surface area per lipid headgroup, directly reducing local lipid packing density (*15, 16*). When restricted by LFA-1 substrate adhesion, this process establishes a stable, compartmentalised gradient of high PM fluidity concentrated at the leading edge. By lowering the local mechanical resistance and bending rigidity of the bilayer interface, this fluidised zone structurally permits the underlying actin cytoskeleton to extend rapidly by enhancing Brownian ratchet-driven actin filament growth at the leading edge. This model proposes that the PM serves as an active, tuneable biophysical gatekeeper that directly coordinates cytoskeletal force generation to drive forward migration.

To further test this requirement for substrate-mediated tension compartmentalisation, we examined the mechanical behaviour of cells on non-adhesive PEG surfaces. We observe a mechanical distinction between the two non-adherent states (Fig 2A, B). In suspension (Fig 2A), cells remain spherical and unconstrained, meaning internal hydrostatic pressure is evenly distributed along the entire cell. Conversely, on PEG under agarose (Fig 2B), cells are physically confined to a 2D plane but cannot engage LFA-1. Notably, the non-adherent condition does not trigger the localised membrane fluidity gradient observed during adhesion on ICAM-1 (Fig 2D). We propose that LFA-1/ICAM-1 engagement focuses hydrostatic pressure locally at the leading edge. Adhesion immobilises a local segment of the actin cortex: consequently, directional water influx drives a local pressure spike, generating mechanical forces that shift the plasma membrane relative to its underlying cytoskeletal anchors (6, 18). This aligns with previous work demonstrating that membrane tension gradients, which are normally equilibrated rapidly across free bilayers, can become restricted and highly localised exclusively within discrete adhesion zones (*24*). Consistent with this, our model is supported by the restriction of the fluidity gradient to the front third of the cell (Fig 2D).

We identified a difference between adhesion-dependent fluidisation required for migration and how actin dynamics are regulated on non-adhesive surfaces. On PEG, T cells cannot locally focus osmotic pressure as effectively as cells on ICAM-1, meaning they lack a distinct fluidity gradient along the cell length (Fig 2B). However, cells on PEG still decrease their global GP slightly (Fig 2B) while maintaining high rates of directional actin retrograde flow (Fig 4G), demonstrating that water flux and cell polarisation can still occur independent of adhesion. Without the anchoring effect of LFA-1 to pin the lipid bilayer and restrict lateral lipid mobility, any localised reduction in lipid packing generated at the leading edge could rapidly equilibrate via lateral diffusion across the cell body. This phenomenon mirrors established models showing that membrane-cytoskeleton attachments act as significant barriers to tension propagation, meaning their absence allows mechanical states to instantly equilibrate out across the unanchored membrane (*24*).

This principle of unconstrained membrane equilibration is further highlighted by our CA rescue experiments. Because externally applied CA incorporates across the entire PM, it fluidises the PM globally rather than reconstructing the leading edge-focused fluidity gradient observed in migrating cells. This subsequent global reduction mechanical resistance is enough to permit rearward movement of the underlying actin engine (Fig 4G), but it only partially rescues forward migration speed (Fig 3E), which requires strict spatial biophysical asymmetry. This phenotypic dissociation reveals a fundamental spatial hierarchy in T cell migration: while actin polymerisation requires only a local reduction in membrane resistance to drive filament elongation, forward cell movement relies on a polarised biophysical gradient to orient that force. Thus, we suggest that while WNK1-dependent water influx acts as the primary driver to lower membrane resistance and allow directional actin assembly, LFA-1-mediated substrate adhesion is required to mechanically compartmentalise fluidisation, preventing PM equilibration to directly convert actin dynamics into directional T cell migration.

Having established that spatial membrane compartmentalisation facilitates cell migration, we next sought to define the biophysical parameters controlling membrane order. Our finding that exposing cells to a hypotonic environment increases PM fluidity, while a hypertonic environment decreases it, demonstrates that osmotic volume changes directly alter lipid packing density (Fig 4C). Disassembling the cortical actin cytoskeleton using latrunculin B increases membrane fluidity to a ‘fluidity plateau’ at a GP of approximately 0.1 that remains unaffected by further hypotonic shock (Fig 4C). This lack of additional fluidisation in a hypotonic environment mirrors the unexpected observation that actin-inhibited cells fail to expand in this condition (Fig 4B). Without the actin cytoskeleton, cells are likely unable to push against the agarose confinement, preventing structured swelling (*30*). Our finding that actin cortical disassembly directly tunes PM order aligns with recent work demonstrating that actin-inhibiting drugs alter PM lipid packing in human immune cells (*31*).

Because the cortical cytoskeleton maintains this tightly constrained baseline, artificially pushing membrane fluidity beyond physiological levels can be detrimental. For example, external CA addition to the PM in vehicle-treated T cells reduced their speed (Fig 3D). Lipid membranes are a finely tuned structure that is optimised to perform required cellular tasks, and the addition of high levels of CA can thus destabilise this structure. Indeed, excessive concentrations of CA have been shown to cause cell death (*32*). This highlights that T cells operate within a highly optimised window of PM fluidity, and disrupting this balance has negative consequences for T cell function.

Beyond the risks of over-fluidisation, the delicate biophysical balance also explains why CA does not fully rescue cell migration speeds in volume-restricted cells, even at increasing concentrations (Fig 3E). It is possible that despite artificial PM fluidisation with CA, the reduced cell size results an altered lipid packing arrangement that physically restricts protein diffusion and prevents the migration machinery from functioning effectively. Alternatively, a reduction in swelling would prevent full untethering of the actin cytoskeleton from the leading edge PM, meaning it remains linked to the membrane and cannot polymerise as effectively. Finally, in the absence of WNK1-dependent swelling, consequent molecular crowding could result in a more viscous cytoplasm (*33, 34*), thereby preventing the efficient diffusion and transport of actin polymerisation machinery.

Despite these physiological limits in volume-restricted cells, establishing this swelling-fluidisation coupling provides a general framework for how cells mechanically regulate lipid order states. This has implications for many cell types, including other immune cells like neutrophils that also swell in response to chemokines (*9*). Further work has shown a critical role for PM properties in regulating immune cell activities including migration and killing (*35*) as well as activation (*36*). Beyond immune cells, cancer cells can similarly rely on water and ion influx to migrate (*37*–*40*). It is thus possible that following osmotic shifts in cancer cells, tumour cell volume and shape changes are enough to initiate migration by locally facilitating actin dynamics due to PM fluidisation. This mechanical shift represents a much simpler mechanism to initiate metastasis compared to larger-scale changes in protein translation, signalling, or lipid synthesis shifts. Indeed, evidence is growing that tumour cells with higher PM fluidity are more likely to metastasise (39), and our work provides evidence for an additional biophysical mechanism that could result in this increased PM fluidity.

Furthermore, this mechanism could be clinically relevant for solid tumour immunology. The environment within solid tumours is often necrotic and consequently rich in ions, resulting in a hypertonic microenvironment (*41*). This hypertonicity would cause cell shrinkage, altered membrane lipid packing and potentially replicates the phenotype of hypertonic-treated cells (Fig 4C), or by extension, WNK463-treated cells (Fig 3B, 5C). Treating engineered T cells, such as CAR T cells, with fluidising agents like CA could increase the biomechanical fitness of these cells in these challenging environments. This would optimise their ability to infiltrate the dense tumour mass, migrate effectively, and perform essential immune functions like cytotoxicity.

In summary, we provide evidence that upon chemokine stimulation, T cells locally reorganise their plasma membrane biophysical properties. This reduction in local lipid packing is functionally coupled to the underlying cytoskeleton, showing that modulating PM fluidity is sufficient to facilitate actin dynamics and thus drive forward migration.

## Materials and methods

### T cell purification and culture

Naïve human CD8^+^ T cells were isolated from peripheral blood mononuclear cells (PBMCs) extracted from buffy coats from healthy donors. Naïve CD8^+^ T cells were isolated using an EasySep Human Naïve CD8^+^ T cell Isolation Kit II (17968, STEMCELL Technologies) according to the manufacturer’s instructions. Following isolation, T cells were rested overnight at 37 °C and 5% CO2 at 1 × 10^6^ cells/ml in RPMI 1640 (Sigma) supplemented with 10% foetal calf serum (FCS) (Sigma), supplemented with 1 mM sodium pyruvate (Gibco) and 20 mM HEPES (Gibco).

### T cell migration under agarose and GP measurements

8-well glass chamber µ-slides (ibidi, 80827) were coated with recombinant human ICAM-1/CD54 Fc chimera (720-IC) at 3 µg/mL in PBS overnight at 4°C. Wells were washed 3x with PBS, blocked for 20 min with 2% bovine serum albumin (Sigma-Aldrich) in PBS, and washed again 3x. For experiments on PEG, 8-well µ-slides were cleaned using 1M HCl for 1h, then washed 3x using MilliQ water, and dried using a nitrogen stream. Slides were plasma activated using O_2_ plasma for 30 sec (CUTE, Femto Science) and immediately coated with 0.2 mg/mL poly(L-lysine)-graft-PEG (SuSoS) in PBS overnight at 4°C. The next day, the slide was washed 3x with PBS.

Agarose gels were adapted from a previously published method (*7*). 0.5% agarose (UltraPure Agarose, ThermoFisher) gels with and RPMI + 10% lipid-depleted FCS (Neuromics) equivalent were prepared at 50°C. Human CCL19 (250 ng/mL, RnD, 361-MI), WNK463 (MedChemExpress, HY-100626) and cholic acid (Larodan, 65-1001, in 96% EtOH) or their equivalent vehicle controls (DMSO, ThermoFisher and 96% EtOH) were added as indicated. To label the plasma membrane, NR12A (*42*) was added to the gel at 400 nM. Gels were left to solidify at room temperature, then cooled to 4°C for 1 h, and subsequently warmed to 37°C and 5% CO_2_ for at least 30 min.

T cells were resuspended in a small volume of RPMI + 2% lipid-depleted FBS and injected under agarose with a pipette. Cells were acclimatised at the microscope for at least 15 min at 37°C. Cells were imaged every 15 s for 5-20 min using either a Zeiss LSM980 or 780 with a 20x (0.8 NA) objective. Images were captured at 1024x1024 resolution in 16 bit using either Zeiss Zen Blue or Black. For NR12A, cells were excited using a 514 laser below 1% power and emission was captured simultaneously between 589-595 nm and 640-645 nm.

To determine the generalised polarisation (GP) metric, images were processed using the Fluidity Calculator FIJI plugin (*43*). Cell shapes were generated from a binary image of the GP signal using the ‘Huang’ auto-threshold algorithm. Resulting images were despeckled, cell shapes were dilated and the ‘fill holes’ binary function was applied. The GP channel and cell shapes were merged into a 2-channel image. To quantify cell migration and GP, the FIJI plugin TrackMate (*44*) was used on the 2-channel image. Dead cells (as determined by average GP <0) as well as tracks shorter than half the video length were excluded. Dead cells (GP < 0) were also excluded.

To determine the GP along the length of the cell, cells were segmented into 8 equally sized segments along the entire width and length of the cell using FIJI.

*Note:* Different optical configurations, detector efficiencies, and acquisition settings between discrete confocal systems introduce baseline offsets in absolute GP values. However, because GP is a ratiometric parameter 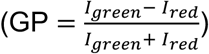, intra-experimental comparative trends remain robust across instruments (*43*).

### Tonicity changes under agarose

8-well glass ibidi µ-slides were washed 2x with PBS. Agarose gel composition was changed by adding, as a fraction of the total volume, either ¼ 2x HBSS (isotonic, 300 mOsm/L), ¼ PBS supplemented with 600 mOsm/L D-mannitol (Sigma, M4125) (hypertonic, 375 mOsm/L), or ¼ MilliQ water (Sigma) (hypotonic, 150 mOsm/L) were prepared according to the recipe described previously. Gels were supplemented with 400 nM NR12A and either latrunculin B (5 µM) or DMSO control. Cells were injected under agarose and immediately imaged on a Zeiss LSM980 using a 20x (0.8 NA) air objective at 37°C. GP was calculated as described previously. Cell outlines (perimeter) and GP were determined using the spot analysis tool in Trackmate.

### Actin retrograde flow measurements

8-well glass ibidi µ-slides were PEG-coated as described previously. Agarose gels were prepared as described previously, with the addition of 1:1000 SPY650-FastAct_X (Spirochrome) prepared in DMSO according to the manufacturer’s instruction. T cells were isolated, rested overnight and stained in 1:1000 SPY650-FastAct_X in T cell medium for 1h at 37°C and 5% CO_2_.

T cells were *not* washed and injected under agarose and visualised 15 min after injection and incubation at 37°C at the microscope. Cells were imaged using a Zeiss LSM980 using a 63x oil objective. Frames were captured every 1 sec for 1 min, proximal to the coverslip. Actin flow rates were determined using kymographs generated in FIJI.

### Flow cytometry stimulation

To measure changes in GP in suspension cells, 30,000 T cells per condition were resuspended in phenol red-free RPMI containing 2% lipid-depleted FCS. Cells were stimulated with either vehicle, CCL19 (250 ng/mL) or α-CD3 and α-CD28 (1 µg/mL each, CD3 OKT3 and CD28.2, both ProteinTech) for 10 min. Cells were subsequently stained with 400 nM NR12A and acquired on a spectral flow cytometer (Cytek Aurora). Flow cytometry data were analysed using a previously described workflow (*31*). Briefly, events were sequentially gated to exclude debris, doublets, and non-viable cells from scattering using FCS Express. GP values were quantified from the fluorescence intensities recorded in the YG2 and YG4 spectral channels using the standard GP equation. Further, dead cells and cells with internalised probe were removed using GP values. Final gating was used to extract GP values for each cell measured and outliers were removed using the ROUT method with a Q value of 1%.

### Cell volume measurement

To measure femtolitre cell volume, T cells were rested overnight. The next day, inhibitor or vehicle control, and/or cholic acid were added for 20 min, and cells were stimulated with either CCL19 (250 ng/mL) or medium for 20 min. Cells were placed in CASYton fluid (Omni Life Science) and median cell volumes were measured using a CASY1 cell counter (Roche).

### Statistical analysis

To calculate P values between two unpaired groups a Mann-Whitney U test was used. To calculate P values when 3 or more groups were compared, we used a 1-way ANOVA or a Kruskal-Wallis test when the data was not normally distributed, as determined by the Shapiro-Wilk test. For experiments when multi-variable conditions are compared (i.e. condition and segment along cell), a 2-way ANOVA was used. In some cases where intracellular measurements were performed, individual cells were used as replicates (time course changes in GP, quantifying GP at the leading edge). All statistical analysis was performed using Prism (GraphPad). Data points are shaped by biological sample (with distinct shapes representing independent experiments).

### Ethics statement

All experiments with human donor samples were approved by the Swedish Ethical Review Authority (Drn 2025-01255-01). All buffy coats were obtained from the Karolinska University Laboratory.

## Acknowledgements

We thank Milka Doktorova and Sezgin lab members for helpful discussion. We acknowledge Arthur Champain for help with optimisation. We thank Franziska Ragaller and Luca Andronico for critical reading of the manuscript. We would also like to thank the Advanced Light Microscopy facility at SciLifeLab for their support with imaging. We thank the donors for their support in providing blood for research purposes. Finally, we thank the funders for their support in making this research possible.

E.S. is supported by Swedish Research Council Grants (grant no. 2020-02682, 2024-02993 and 2024-00289), Wellcome Leap’s Dynamic Resilience Program (jointly funded by Temasek Trust), CancerFonden (25 4592 Pj) Karolinska Institutet (2024-03250; 2024-03341; 2022-00803; 2020-00997), Cancer Research KI (2024-03488), Strategic Research Programme in Diabetes at Karolinska Institutet, Human Frontier Science Program (RGP0025/2022), Longevity Impetus Grant from Norn Group, Hevolution Foundation and Rosenkranz Foundation. E.S. is an EMBO Young Investigator (YIP-2025). L.L.d.B. is supported by a SciLifeLab Research Environment Development postdoctoral fellowship.

## Author contributions

Conceptualisation: L.L.d.B., E.S.

Data curation: L.L.d.B.

Formal analysis: L.L.d.B., C.O.G.

Funding acquisition: E.S., L.L.d.B.

Investigation: L.L.d.B.

Methodology: L.L.d.B., C.O.G., E.S.

Project administration: L.L.d.B., E.S.

Resources: L.L.d.B., E.S.

Supervision: E.S.

Visualisation: L.L.d.B.

Writing: L.L.d.B., E.S.

## Supporting Movie Legends

**Supporting Movie 1:** Migration of stimulated and unstimulated human CD8+ T cells

**Supporting Movie 2:** Tracking of single cells and quantification of membrane fluidity during migration

**Supporting Movie 3:** Retrograde actin flow for stimulated cells with WNK463 and cholic acid treatment.

## References

1. M. F. Krummel, F. Bartumeus, A. Gérard, T cell migration, search strategies and mechanisms. Nat. Rev. Immunol. 16, 193–201 (2016).

2. M. Huse, Mechanical forces in the immune system. Nat. Rev. Immunol. 17, 679–690 (2017).

3. S. B. Thompson, M. M. Waldman, J. Jacobelli, Polymerization power: effectors of actin polymerization as regulators of T lymphocyte migration through complex environments. FEBS J., febs.16130 (2021).

4. D. Démoulin, M. F. Carlier, J. Bibette, J. Baudry, Power transduction of actin filaments ratcheting in vitro against a load. Proc. Natl. Acad. Sci. U. S. A. 111, 17845–17850 (2014).

5. C. S. Peskin, G. M. Odell, G. F. Oster, Cellular motions and thermal fluctuations: the Brownian ratchet. Biophys. J. 65, 316–324 (1993).

6. R. Köchl, F. Thelen, L. Vanes, T. F. Brazão, K. Fountain, J. Xie, C.-L. Huang, R. Lyck, J. V. Stein, V. L. J. Tybulewicz, WNK1 kinase balances T cell adhesion versus migration in vivo. Nat. Immunol. 17, 1075–1083 (2016).

7. L. L. de Boer, L. Vanes, S. Melgrati, J. Biggs O’May, D. Hayward, P. C. Driscoll, J. Day, A. Griffiths, R. Magueta, A. Morrell, J. I. MacRae, R. Köchl, V. L. J. Tybulewicz, T cell migration requires ion and water influx to regulate actin polymerization. Nat. Commun. 14, 7844 (2023).

8. J. Biggs O’May, L. Vanes, L. L. de Boer, D. A. Lewis, H. Hartweger, S. Kunzelmann, D. Hayward, M. Llorian, R. Köchl, V. L. J. Tybulewicz, WNK1-dependent water influx is required for CD4+ T cell activation and T cell-dependent antibody responses. Nat. Commun. 16, 1857 (2025).

9. T. L. Nagy, E. Strickland, O. D. Weiner, Neutrophils actively swell to potentiate rapid migration. eLife 12, RP90551 (2024).

10. T. Karlsson, B. C. Lagerholm, E. Vikström, V. M. Loitto, K. E. Magnusson, Water fluxes through aquaporin-9 prime epithelial cells for rapid wound healing. Biochem. Biophys. Res. Commun. 430, 993–998 (2013).

11. T. Karlsson, A. Bolshakova, M. A. Magalhães, V. M. Loitto, K. E. Magnusson, Fluxes of Water through Aquaporin 9 Weaken Membrane-Cytoskeleton Anchorage and Promote Formation of Membrane Protrusions. PLoS ONE 8 (2013).

12. L. Miguel, D. M. Owen, C. Lim, C. Liebig, J. Evans, A. I. Magee, E. C. Jury, Primary Human CD4+ T Cells Have Diverse Levels of Membrane Lipid Order That Correlate with Their Function. J. Immunol. 186, 3505–3516 (2011).

13. P. Sens, M. S. Turner, Budded membrane microdomains as tension regulators. Phys. Rev. E 73, 031918 (2006).

14. L. Guillou, A. Babataheri, M. Saitakis, A. Bohineust, S. Dogniaux, C. Hivroz, A. I. Barakat, J. Husson, P. Bassereau, T-lymphocyte passive deformation is controlled by unfolding of membrane surface reservoirs. Mol. Biol. Cell 27, 3574–3582 (2016).

15. D. Marsh, Lateral Pressure Profile, Spontaneous Curvature Frustration, and the Incorporation and Conformation of Proteins in Membranes. Biophys. J. 93, 3884–3899 (2007).

16. A. Hossein, M. Deserno, Spontaneous Curvature, Differential Stress, and Bending Modulus of Asymmetric Lipid Membranes. Biophys. J. 118, 624–642 (2020).

17. E. S. Welf, C. E. Miles, J. Huh, E. Sapoznik, J. Chi, M. K. Driscoll, T. Isogai, J. Noh, A. D. Weems, T. Pohlkamp, K. Dean, R. Fiolka, A. Mogilner, G. Danuser, Actin-Membrane Release Initiates Cell Protrusions. Dev. Cell 55, 723–736.e8 (2020).

18. A. Bisaria, A. Hayer, D. Garbett, D. Cohen, T. Meyer, Membrane-proximal F-actin restricts local membrane protrusions and directs cell migration. Science 368, 1205–1210 (2020).

19. E. Moeendarbary, L. Valon, M. Fritzsche, A. R. Harris, D. A. Moulding, A. J. Thrasher, E. Stride, L. Mahadevan, G. T. Charras, The cytoplasm of living cells behaves as a poroelastic material. Nat. Mater. 12, 253–261 (2013).

20. G. T. Charras, M. Coughlin, T. J. Mitchison, L. Mahadevan, Life and Times of a Cellular Bleb. Biophys. J. 94, 1836–1853 (2008).

21. F. Heinemann, S. K. Vogel, P. Schwille, Lateral Membrane Diffusion Modulated by a Minimal Actin Cortex. Biophys. J. 104, 1465–1475 (2013).

22. F. Schneider, D. Waithe, M. P. Clausen, S. Galiani, T. Koller, G. Ozhan, C. Eggeling, E. Sezgin, R. G. Parton, Diffusion of lipids and GPI-anchored proteins in actin-free plasma membrane vesicles measured by STED-FCS. Mol. Biol. Cell 28, 1507–1518 (2017).

23. E. Sezgin, I. Levental, S. Mayor, C. Eggeling, The mystery of membrane organization: composition, regulation and roles of lipid rafts. Nat. Rev. Mol. Cell Biol. 18, 361–374 (2017).

24. J. M. García-Arcos, A. Mehidi, J. Sanchez-Velasquez, P. Guillamat, C. Tomba, L. Houzet, L. Capolupo, J. Espadas, G. D’Angelo, A. Colom, E. Hinde, C. Aumeier Roux, Adherent cells sustain membrane tension gradients independently of migration. Nat. Commun. 16, 10539 (2025).

25. D. I. Danylchuk, S. Moon, K. Xu, A. S. Klymchenko, Switchable Solvatochromic Probes for Live-Cell Super-resolution Imaging of Plasma Membrane Organization. Angew. Chem. 131, 15062–15066 (2019).

26. M. Hons, A. Kopf, R. Hauschild, A. Leithner, F. Gaertner, J. Abe, J. Renkawitz, J. V. Stein, M. Sixt, Chemokines and integrins independently tune actin flow and substrate friction during intranodal migration of T cells. Nat. Immunol. 19, 606–616 (2018).

27. K. Yamada, H.-M. M. Park, D. F. Rigel, K. DiPetrillo, E. J. Whalen, A. Anisowicz, M. Beil, J. Berstler, C. E. Brocklehurst, D. A. Burdick, S. L. Caplan, M. P. Capparelli, G. Chen, W. Chen, B. Dale, L. Deng, F. Fu, N. Hamamatsu, K. Harasaki, T. Herr, P. Hoffmann, Q.-Y. Y. Hu, W.-J. J. Huang, N. Idamakanti, H. Imase, Y. Iwaki, M. Jain, J. Jeyaseelan, M. Kato, V. K. Kaushik, D. Kohls, V. Kunjathoor, D. LaSala, J. Lee, J. Liu, Y. Luo, F. Ma, R. Mo, S. Mowbray, M. Mogi, F. Ossola, P. Pandey, S. J. Patel, S. Raghavan, B. Salem, Y. H. Shanado, G. M. Trakshel, G. Turner, H. Wakai, C. Wang, S. Weldon, J. B. Wielicki, X. Xie, L. Xu, Y. I. Yagi, K. Yasoshima, J. Yin, D. Yowe, J.-H. H. Zhang, G. Zheng, L. Monovich, Small-molecule WNK inhibition regulates cardiovascular and renal function. Nat. Chem. Biol. 12, 896–898 (2016).

28. M. C. Neves, H. A. L. Filipe, R. L. Reis, J. P. Prates Ramalho, F. Coreta-Gomes, M. J. Moreno, L. M. S. Loura, Interaction of Bile Salts With Lipid Bilayers: An Atomistic Molecular Dynamics Study. Front. Physiol. 10 (2019).

29. D. Holz, D. Vavylonis, Building a dendritic actin filament network branch by branch: models of filament orientation pattern and force generation in lamellipodia. Biophys. Rev. 10, 1577–1585 (2018).

30. G.-X. Wang, Y.-P. Dai, S. Bongalon, W. J. Hatton, K. Murray, J. R. Hume, I. A. Yamboliev, Hypotonic Activation of Volume-sensitive Outwardly Rectifying Anion Channels (VSOACs) Requires Coordinated Remodeling of Subcortical and Perinuclear Actin Filaments. J. Membr. Biol. 208, 15–26 (2005).

31. C. O. Gurdap, D. Aydos, L. A. Andronico, G. Tóth, T. Ceker, J. Cowgill, I. M. Akbulut Koyuncu, N. Basak, J. Mikes, A. S. Klymchenko, F. Pietrocola, P. Brodin, I. Lanekoff, V. C. Bitirim, E. Sezgin, Spectral biophysical cytometry with nanosensors reveals remodelling of immune cells in atherosclerosis. Nat. Nanotechnol., 1–8 (2026).

32. Y. Zhou, K. N. Maxwell, E. Sezgin, M. Lu, H. Liang, J. F. Hancock, E. J. Dial, L. M. Lichtenberger, I. Levental, Bile Acids Modulate Signaling by Functional Perturbation of Plasma Membrane Domains. J. Biol. Chem. 288, 35660–35670 (2013).

33. A. Miermont, F. Waharte, S. Hu, M. N. McClean, S. Bottani, S. Léon, P. Hersen, Severe osmotic compression triggers a slowdown of intracellular signaling, which can be explained by molecular crowding. Proc. Natl. Acad. Sci. U. S. A. 110, 5725–5730 (2013).

34. P. Nunes, I. Roth, P. Meda, E. Féraille, D. Brown, U. Hasler, Ionic imbalance, in addition to molecular crowding, abates cytoskeletal dynamics and vesicle motility during hypertonic stress. Proc. Natl. Acad. Sci. 112, E3104–E3113 (2015).

35. L. A. Andronico, C. O. Gurdap, A. Arora, F. Ragaller, P. A. Sandoz, Y. Jiang, S. Giatrellis, L. L. de Boer, V. Carannante, S. Iskrak, J. Mikes, M. Buggert, A. Österborg, B. Önfelt, A. S. Klymchenko, P. Brodin, E. Sezgin, Plasma membrane order maps functional diversity in immune cells. Nat. Chem. Biol., 1–12 (2026).

36. J. Dinic, A. Riehl, J. Adler, I. Parmryd, The T cell receptor resides in ordered plasma membrane nanodomains that aggregate upon patching of the receptor. Sci. Rep. 5, 10082 (2015).

37. K. M. Stroka, H. Jiang, S. H. Chen, Z. Tong, D. Wirtz, S. X. Sun, K. Konstantopoulos, Water permeation drives tumor cell migration in confined microenvironments. Cell 157, 611–623 (2014).

38. T. Garzon-Muvdi, P. Schiapparelli, C. apRhys, H. Guerrero-Cazares, C. Smith, D. H. Kim, L. Kone, H. Farber, D. Y. Lee, S. S. An, A. Levchenko, A. Quiñones-Hinojosa, Regulation of brain tumor dispersal by NKCC1 through a novel role in focal adhesion regulation. PLoS Biol. 10, e1001320 (2012).

39. P. Schiapparelli, H. Guerrero-Cazares, R. Magaña-Maldonado, S. M. Hamilla, S. Ganaha, E. Goulin Lippi Fernandes, C. H. Huang, H. Aranda-Espinoza, P. Devreotes, A. Quinones-Hinojosa, NKCC1 Regulates Migration Ability of Glioblastoma Cells by Modulation of Actin Dynamics and Interacting with Cofilin. EBioMedicine 21, 94–103 (2017).

40. A. B. Jaykumar, J. U. Jung, P. K. Parida, T. T. Dang, C. Wichaidit, A. R. Kannangara, S. Earnest, E. J. Goldsmith, G. W. Pearson, S. Malladi, M. H. Cobb, WNK1 Enhances Migration and Invasion in Breast Cancer Models. Mol. Cancer Ther. 20, 1800–1808 (2021).

41. R. Eil, S. K. Vodnala, D. Clever, C. A. Klebanoff, M. Sukumar, J. H. Pan, D. C. Palmer, A. Gros, T. N. Yamamoto, S. J. Patel, G. C. Guittard, Z. Yu, V. Carbonaro, K. Okkenhaug, D. S. Schrump, W. M. Linehan, R. Roychoudhuri, N. P. Restifo, Ionic immune suppression within the tumour microenvironment limits T cell effector function. Nature 537, 539–543 (2016).

42. D. I. Danylchuk, S. Moon, K. Xu, A. S. Klymchenko, Switchable Solvatochromic Probes for Live-Cell Super-resolution Imaging of Plasma Membrane Organization. Angew. Chem. 131, 15062–15066 (2019).

43. P. Carravilla, L. Andronico, J. Schlegel, Y. B. Urem, E. Sjule, F. Ragaller, F. Weber, C. O. Gurdap, Y. Ascioglu, T. Sych, J. Lorent, E. Sezgin, Measuring plasma membrane fluidity using confocal microscopy. Nat. Protoc. 20, 1976–2004 (2025).

44. J. Y. Tinevez, N. Perry, J. Schindelin, G. M. Hoopes, G. D. Reynolds, E. Laplantine, S. Y. Bednarek, S. L. Shorte, K. W. Eliceiri, TrackMate: An open and extensible platform for single-particle tracking. Methods 115, 80–90 (2017).

